# Structural analysis of phosphorylation-associated interactions of human MCC to Scribble PDZ domains

**DOI:** 10.1101/501064

**Authors:** Sofia Caria, Bryce Z. Stewart, Patrick O. Humbert, Marc Kvansakul

**Affiliations:** Department of Biochemistry & Genetics, La Trobe Institute for Molecular Science, La Trobe University, Melbourne, Victoria 3086, Australia.; SAXS/WAXS, Australian Synchrotron, 800 Blackburn Road, Clayton, VIC 3168, Australia; Research Centre for Molecular Cancer Prevention, La Trobe University, Melbourne, Victoria 3086, Australia.; Department of Biochemistry & Molecular Biology, University of Melbourne, Melbourne, Victoria 3010, Australia; Department of Clinical Pathology, University of Melbourne, Melbourne, Victoria 3010, Australia

**Keywords:** Scribble, MCC, phosphorylation, PDZ domain, cell polarity, X-ray crystallography

## Abstract

Scribble is a crucial adaptor protein that plays a pivotal role during establishment and control of cell polarity, impacting many physiological processes ranging from cell migration to immunity and organization of tissue architecture. Scribble harbors a leucine-rich repeat domain and four PDZ domains, which mediate most of Scribble’s interactions with other proteins. It has become increasingly clear that posttranslational modifications substantially impact Scribble-ligand interactions, with phosphorylation being a major modulator of binding to Scribble. To better understand how Scribble PDZ domains direct cell polarity signalling and how phosphorylation impacts this process, we investigated human Scribble interactions with MCC (mutated in colorectal cancer). We systematically evaluated the ability of all four individual Scribble PDZ domains to bind the PDZ-binding motif (PBM) of MCC as well as MCC phosphorylated at the -1 Ser position. We show that Scribble PDZ1 and PDZ3 are the major interactors with MCC, and that modifications to Ser at the -1 position in the MCC PBM only has a modest effect on binding to Scribble PDZ domains. We then examined the structural basis for these observations by determining the crystal structures of Scribble PDZ1 domain bound to both the unphosphorylated MCC PBM as well as phosphorylated MCC. Our structures indicated that phospho-Ser at the -1 position in MCC is not involved in major contacts with Scribble PDZ1, and in conjunction with our affinity measurements suggest that the impact of phosphorylation at the -1 position of MCC extends beyond a simple modulation of the affinity for Scribble PDZ domains.

## INTRODUCTION

Cell polarity, which manifests itself as the asymmetric distribution of cellular constituents as well as proteins, lipids and carbohydrates into distinct cellular domains (1), is a critical property of eukaryotic cells and pivotal for correct tissue architecture and tissue development. The particular distribution of cellular constituents leads to the establishment of apical-basal cell polarity in epithelial cells, and impacts numerous critical cellular signaling pathways including those involved in apoptosis, vesicle trafficking, cell proliferation and migration (2). Furthermore, loss of cell polarity is recognized as an important hallmark of cancer development (3), underscoring the importance of correct cell polarity for healthy tissues. Apico-basal polarity is controlled by the interplay of three multi-protein complexes, Par, Crumbs and Scribble, with epithelial cell polarity being orchestrated by the antagonistic interaction between PAR and Crumbs with the Scribble complex. In mammals, the Scribble complex comprises Scribble (SCRIB), one of 4 Dlg (Discs Large) homologs (DLG1-4) and 2 Lgl (Lethal Giant Larvae) homologs (LLGL1, LLGL2) which are highly conserved from the vinegar fly to humans (4). Scribble was identified in *Drosophila melanogaster* as a tumour suppressor where loss of Scribble resulted in disrupted epithelial tissue organization accompanied by aberrant growth in the imaginal discs of the larvae (5). The ability to suppress tumors was shown to be conserved across species, with Scribble knock-out promoting tumor initiation, and when coupled with oncogenic drivers including RAS, tumor progression in diverse epithelial tissues including mammary, prostate, skin and the lung (6-10).

Scribble is a large multi-domain scaffold protein comprising 16 Leucine Rich Repeats and 4 PSD-95/Disc-large/ZO-1 (PDZ) domains, and belongs to the LAP family of proteins. These domains enable Scribble to interact with a diverse set of interactors that play a role in a range of discrete signaling pathways (4). Whilst the majority of interactions are modulated by the four PDZ domains, the Scribble LRR domain also engages a specific subset of interactions, such as with Lgl2, during the regulation of cell polarity (11).

Binding to Scribble via its PDZ domains is typically mediated via C-terminally located PDZ-binding motifs (PBMs) on specific interactors. Although these sequences are specific, several studies have shown that whilst PBMs on Scribble interactors selectively engage Scribble, the Scribble PDZ domains appear to harbor overlapping preferences for certain ligands, with each PDZ domain capable of engaging multiple binding partners. Scribble PDZ domains are classified as Class I PDZ domains, which recognizes a consensus X-T/S-X-Ø_COOH_ motif (where X can be any amino acid residue, and Ø is a hydrophobic residue) in the PBM of interactors. In addition to amino acid variations, post-translational modification via phosphorylation has also been shown to impact ligand binding to PDZ domains. Binding of Kir2.3 (12) and stargazin (13) to DLG4 (PSD-95) is regulated by phosphorylation, with phosphorylation of a Thr at the -2 position in stargazin abrogating binding to DLG4. Similarly, phosphorylation of a Ser at -2 prevents Kir2.3 binding to DLG4. More recently phosphorylation of PBMs was examined using proteomics approaches, which suggested that phosphorylation of PBMs occurs frequently and can have diverse impacts on PDZ domain binding (14). A comprehensive analysis of the effect of phosphorylation in the PBM of R58 in position -2, -3, -5 and -6 on the binding to the SNX27 PDZ domain (15) revealed that phosphorylation of the -2 position blocks the interaction, whereas phosphorylation at the -3 and -5 position significantly increased affinity.

MCC (mutated in colorectal cancer) has been shown to be a Scribble interactor (16). This complex appears to be highly conserved evolutionarily as *Drosophila* Scrib and MCC can interact physically (16) and in zebrafish loss of Scrib and MCC interact genetically to regulate convergent extension movements during gastrulation (17). MCC is a large multi-PDZ domain protein found in epithelial cells that has been shown to participate in several key cellular signaling pathways including canonical and non-canonical Wnt signaling, NFKB signaling as well as cell cycle control (17-20). More recently, it was shown that MCC harbors a C-terminal PBM that is able to engage human Scribble PDZ1 and 3 domains to impact epithelial cell polarity (16). Interestingly, the MCC PBM has been shown to be a target for phosphorylation in position -1 which features a Ser residue, with phosphorylation impacting the formation of lamellipodia in colon epithelial cells (21). Apart from MCC only Syndecan1 has been described to feature phosphorylation a the -1 position of their PBM (22), which promoted binding to the Tiam1 PDZ domain by improving the affinity from 51.8 to 35.1 μM.

Notably, phosphorylation of the -1 Ser in the MCC PBM was proposed to increase binding to Scribble, as evidenced by increased immunoprecipitation of phospho-MCC. The impact on phosphorylation of the MCC PBM on binding to the Scribble PDZ1 domain was recently shown to lead to a nearly two-fold increase in affinity from 4.4 μM for MCC to 2.4 μM for phospho-MCC as measured by microscale thermophoresis and ITC (14), however NMR chemical shift analysis revealed no significant differences in binding to the Scribble PDZ1 for unphosphorylated and phosphorylated MCC (23).

To understand the impact of phosphorylation on the ability of the MCC PBM to bind to Scribble PDZ domains, we systematically examined the affinity of recombinant human Scribble PDZ domains for peptides spanning the MCC PBM with either phosphorylated or unphosphorylated Ser at the -1 position, as well as peptides harboring a phosphomimic Ser to Glu mutation. Finally, we determined crystal structures of human Scribble PDZ1 bound to a MCC PBM peptide as well as the MCC PBM with phosphorylated Ser at the -1 position to examine the structural basis for this interaction and the impact of phosphorylation on binding to Scribble PDZ1. Our findings suggest that phosphorylation of the MCC PBM at the -1 position is not a major determinant of interactions with human Scribble PDZ1 and 3 domains.

## RESULTS

### Isolated human Scribble PDZ domains specifically interact with the β-PIX PBM

Human Scribble has previously been shown to directly interact with the MCC C-terminal PDZ binding motif (PBM) via its PDZ1 and 3 domains using pull-down and biochemical assays. To understand the impact of phosphorylation on the binding of MCC to Scribble we examined the affinity of recombinant human Scribble PDZ1, 2, 3 and 4 domains for a panel of 8-mer peptide corresponding to the MCC PBM (Figure 1, Table 1). Our ITC measurements revealed that unphosphorylated MCC bound to Scribble PDZ1 and 3 domains with K_D_ values of 7.7 and 5.0 μM, respectively, and did not show detectable binding to Scribble PDZ2 and 4 domains. A phosphorylated MCC peptide carrying a phosphorylated Ser at the -1 position bound Scribble PDZ1 with 6.1 μM and PDZ3 with 3.7 μM, whereas two phosphomimic MCC peptides with a Ser to Glu (MCC_SE) or a Ser to Asp (MCC_SD) mutation bound PDZ1 with 7.2 or 7.0 μM, whereas PDZ3 was bound with 4.5 or 3.4 μM, respectively (Figure 2, Table 1).

**Table 1:** Summary of affinity and thermodynamic binding parameters for Scrib PDZ domain interactions with MCC peptides. NB denotes no binding. Each of the value was calculated from at least three independent experiments.

| | $K_d$ (nM) | $\Delta H$ (kcal/mol) | $-T\Delta S$<br>(cal/mol/K) | N |
| --- | --- | --- | --- | --- |
| PDZ1:MCC | 7715±499 | -4.53±0.06 | -2.44±0.09 | 1.1±0.05 |
| PDZ2:MCC | NB | NB | NB | NB |
| PDZ3:MCC | 4971±901 | -13.38±0.44 | 6.12±0.45 | 1.1±0.02 |
| PDZ4:MCC | NB | NB | NB | NB |
| PDZ1:pMCC | 6083±555 | -3.70±0.99 | -3.41±0.93 | 1.0±0.032 |
| PDZ2:pMCC | NB | NB | NB | NB |
| PDZ3:pMCC | 3744±788 | -9.99±1.00 | 2.58±1.12 | 1.0±0.03 |
| PDZ4:pMCC | NB | NB | NB | NB |
| PDZ1:MCC_SE | 7185±884 | -3.53±0.22 | -3.49±0.15 | 1.0±0.05 |
| PDZ3:MCC_SE | 4476±510 | -8.14±0.64 | 2.82±0.84 | 1.0±0.05 |
| PDZ1:MCC_SD | 7033±754 | -3.14±0.12 | -3.90±0.13 | 1.0±0.02 |
| PDZ3:MCC_SD | 3441±632 | -8.32±0.79 | 0.86±0.69 | 1.1±0.01 |

**FIGURE 1:**
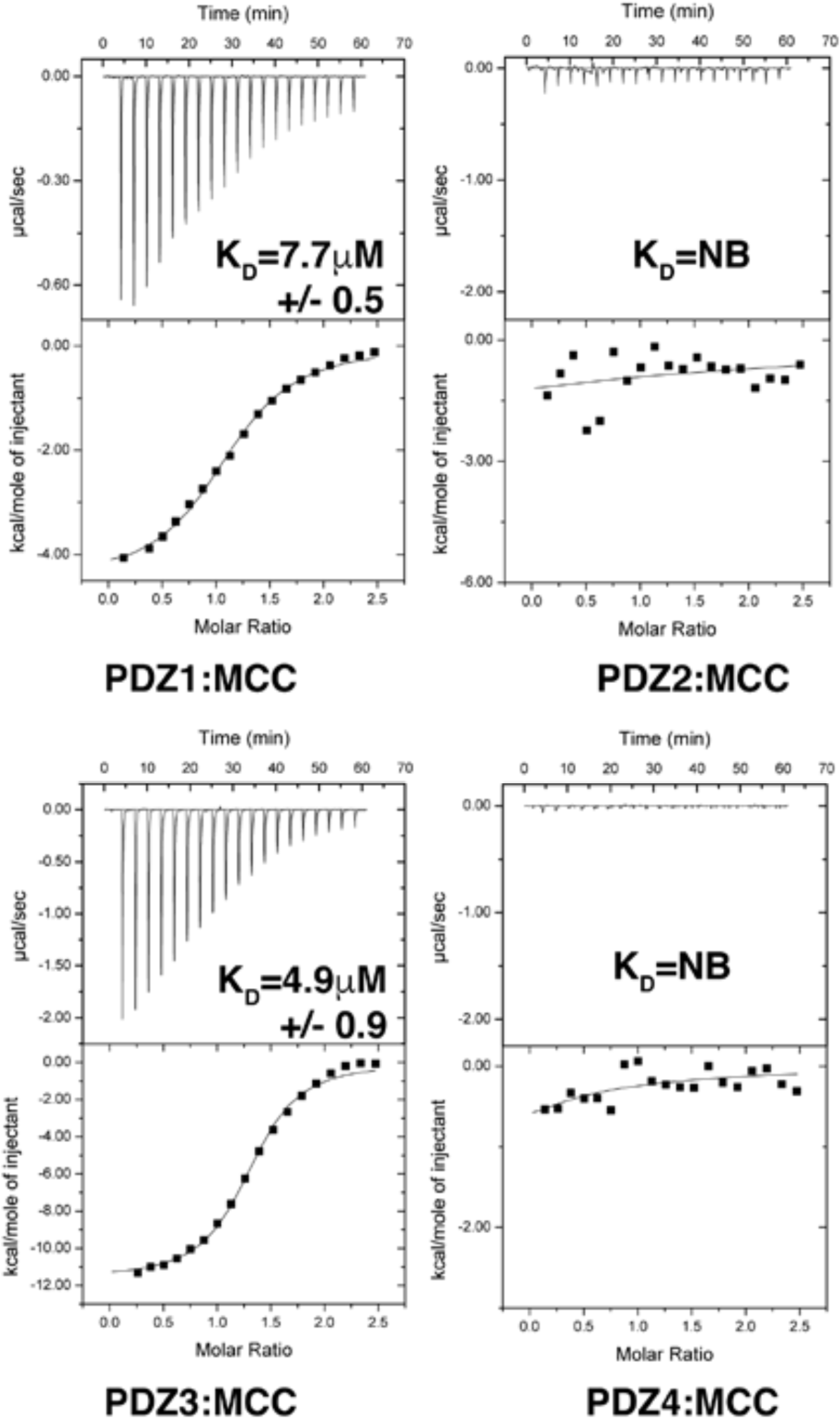
Interaction profiles of Scribble PDZ domains with MCC peptide. Binding profiles of isolated Scribble PDZ domains interaction with MCC peptides are displayed. Each profile is represented by a raw thermogram (top panel) and a binding isotherm fitted with a one-site binding model (bottom panels). K_D_: dissociation constant; ±: standard deviation; NB: no binding. Each of the value was calculated from at least three independent experiments.

**FIGURE 2:**
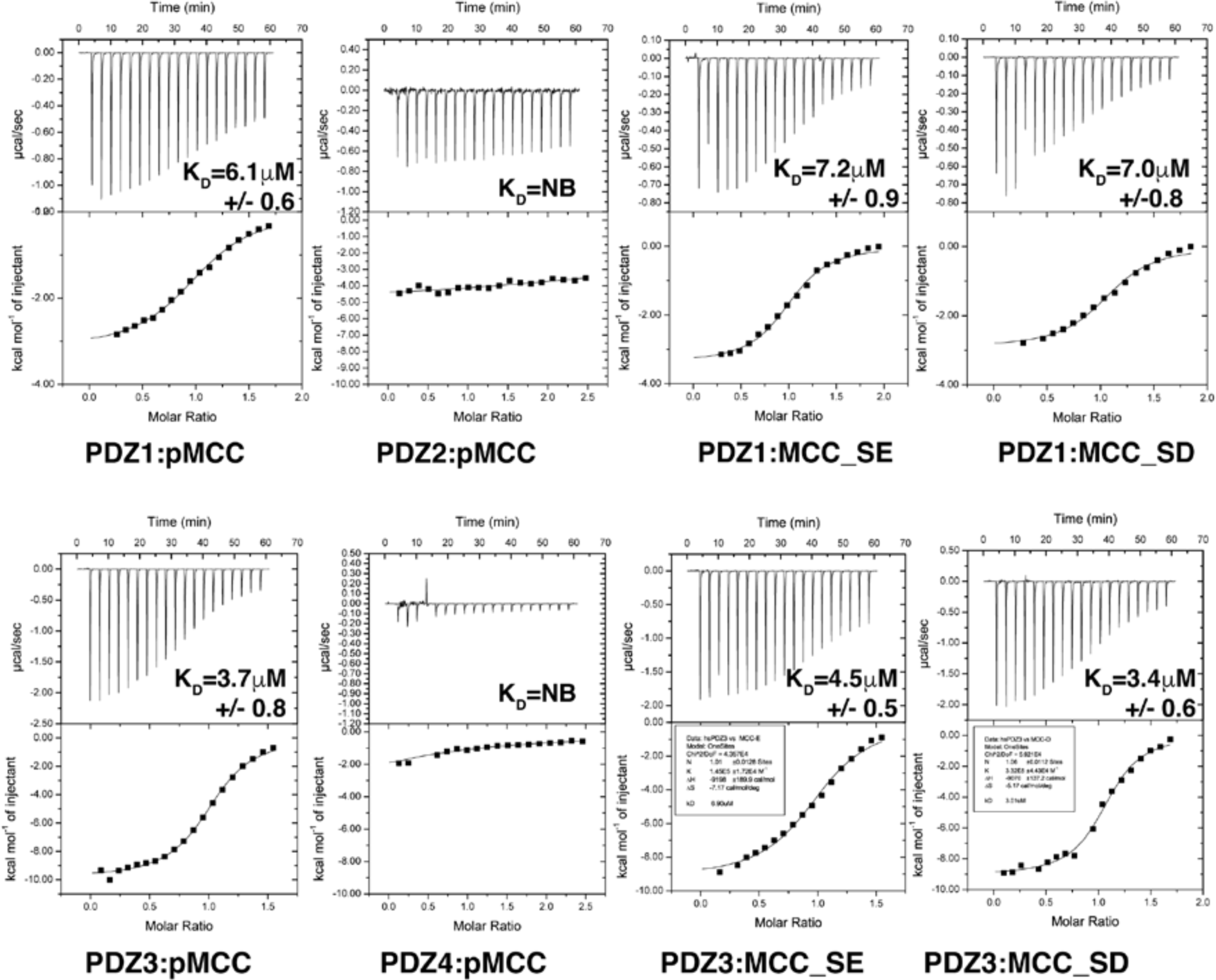
Interaction profiles of Scribble PDZ domains with phosphorylated MCC peptide or phosphomimicking MCC peptides. Isothermal titration calorimetry binding profiles of isolated PDZ domains interaction with phosphorylated MCC (pMCC) peptide or MCC peptides bearing phosphomimicking substitutions. Each profile is represented by a raw thermogram (top panel) and a binding isotherm fitted with a one-site binding model (bottom panels). Each binding profile is a representative example from three independent experiments. K_D_: dissociation constant; ±: standard deviation; NB: no binding. Each of the value was calculated from at least three independent experiments. Peptides used are MCC (PHTNETSL), pMCC (PHTNET(phoS)L), MCC_SE (PHTNETEL) and MCC_SD (PHTNETDL).

Examination of the thermodynamic binding parameters of MCC binding to Scribble PDZ3 indicates that the modestly tighter binding of phosphorylated MCC (pMCC) to Scribble PDZ3 compared to the unphosphorylated MCC peptide is largely driven by a more favorable-TΔS or entropic term, indicative of more disorder upon pMCC binding (Figure 3, Table 1).

**FIGURE 3:**
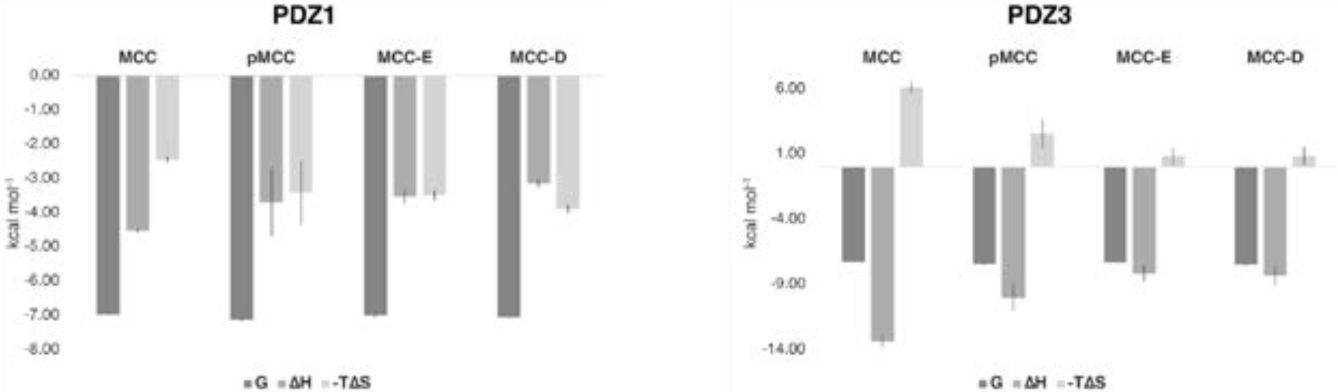
Thermodynamic analysis of Scribble PDZ1 and PDZ3 domain binding to wild-type and modified MCC peptides. The contribution of -ΔH (kcal/mol) and TΔS (cal/mol/K) to the binding of wild type MCC as well as pMCC, MCC_SE and MCC_SD are shown. Each of the value was calculated from at least three independent experiments. Peptides used are pMCC (PHTNET(phoS)L), MCC_SE (PHTNETEL) and MCC_SD (PHTNETDL).

### The crystal structures of SCRIB PDZ1:MCC peptides

To understand the structural implications of Ser phosphorylation at the -1 site of the MCC PBM we next determined the crystal structure of PDZ1 bound to the MCC 8-mer peptide (Figure 4, 5, Table 2). As previously shown (24) PDZ1 adopts a compact globular fold comprising six β-strands and two α-helices that form a β-sandwich structure. The MCC peptide is bound in the canonical ligand binding groove formed by the β2 strand and helix α2. Compared to the previously determined structure of a Scribble PDZ1:β-PIX complex or ligand free Scribble PDZ1 (PDB ID 5VWC) (24), binding of MCC does not significantly alter the PDZ1 domain structure, with the rmsd between Scribble PDZ1:β-PIX and MCC complexes being 1.3 Å over 92 Cα atoms, and between PDZ1 alone and PDZ1:MCC being 0.9 Å over 99 Cα atoms (25).

**Table 2:** Data collection and refinement statistics.

|  | PDZ1:MCC | PDZ1:pMCC |
| --- | --- | --- |
| **Data collection** |  |  |
| Space group | I 121 | I 4<sub>1</sub> |
| No of molecules in AU | 2+2 | 2+2 |
| Cell dimensions |  |  |
| *a*, *b*, *c* (Å) | 57.40, 55.56, 61.78 | 53.60, 53.60, 214.45 |
| $\alpha$, $\beta$, $\gamma$ (°) | 90.00, 116.12, 90.00 | 90.00, 90.00, 90.00 |
| Wavelength (Å) | 0.9537 | 0.9537 |
| Resolution (Å)* | 39.25-2.60 (2.71-2.60) | 4288-2.14 (2.48-2.14) |
| *R*<sub>sym</sub> or *R*<sub>merge</sub> * | 0.085 (0.936) | 0.082 (0.750) |
| *I* / $\sigma I$ * | 7.3 (1.8) | 12.6 (1.5) |
| CC(1/2) | 0.995 (0.573) | 0.999 (0.641) |
| Completeness spherical (%)* | 97.6 (93.8) | 53.3 (8.0) |
| Completeness ellipsoidal (%)* | - | 91.5 (69.1) |
| Redundancy* | 2.8 (2.9) | 5.7 (4.5) |
| Wilson B-factor | 22.02 | 41.65 |
| **Refinement** |  |  |
| Resolution (Å) | 37.79-2.60 | 42.88-2.14 |
| No. reflections | 5311 | 8810 |
| *R*<sub>work</sub> / *R*<sub>free</sub> | 0.223/0.273 | 0.2112/0.2447 |
| No. non-hydrogen atoms |  |  |
| Protein | 1574 | 1611 |
| Ligand/ion | 22 | 75 |
| Water | 4 | 34 |
| *B*-factors |  |  |
| Protein | 64.31 | 43.64 |
| Ligand/ion | 68.43 | 47.07 |
| Water | 60.62 | 38.33 |
| R.m.s. deviations |  |  |
| Bond lengths (Å) | 0.005 | 0.006 |
| Bond angle (°) | 1.00 | 1.22 |
| Ramachandran plot (%) |  |  |
| Favored | 96.95 | 98.5 |
| Allowed | 3.05 | 1.5 |
| Disallowed | 0 | 0 |

\* Data in parentheses are for highest resolution shell

**FIGURE 4:**
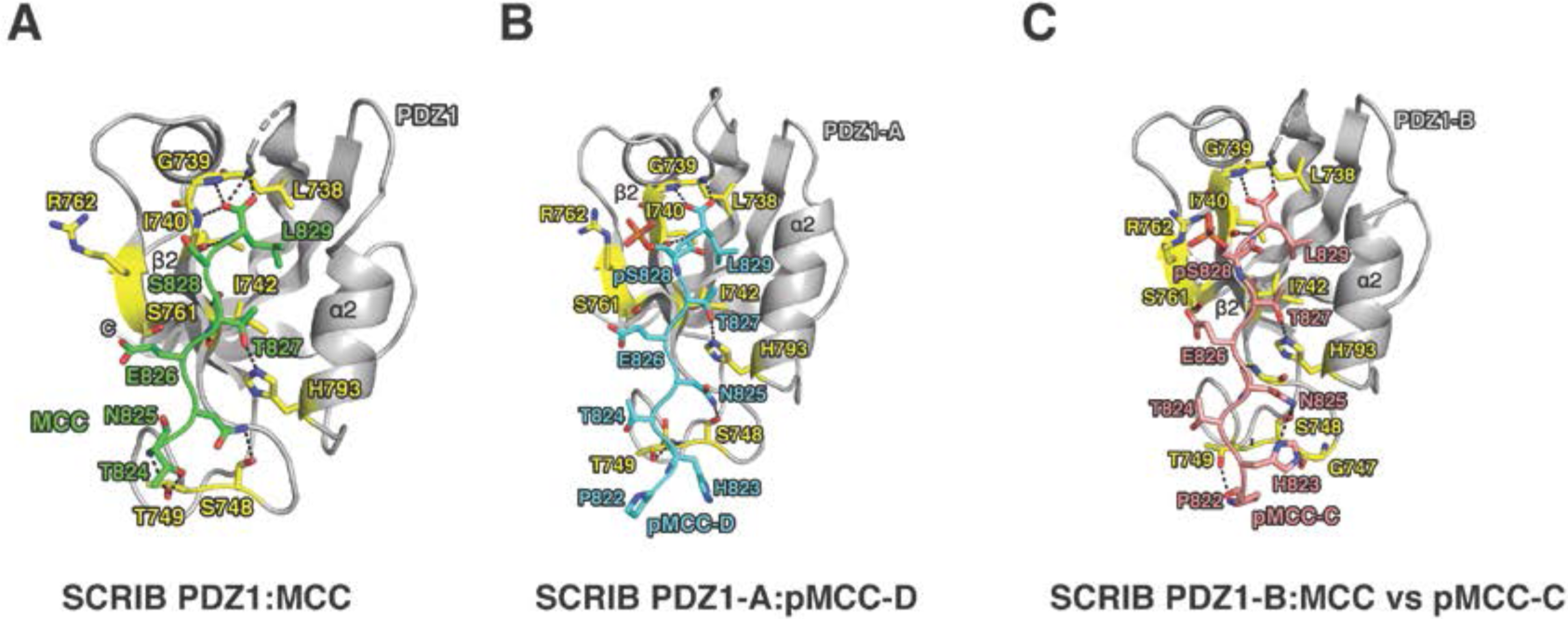
The crystal structures of PDZ1 bound to wild-type MCC and phosphorylated MCC peptide. The MCC peptide engages the Scribble PDZ1 domain via a shallow groove located between the β2 and α2. (A) PDZ1 (grey and yellow) is shown as a cartoon with unphosphorylated MCC peptide (green) represented as sticks. (B,C) PDZ1 (grey and yellow) is shown as a cartoon with phosphorylated MCC peptide (cyan or coral) represented as sticks. Both copies of the complexes formed by PDZ1 chain A with pMCC chainD as well as PDZ1 chain B and pMCC chain C are shown.

**FIGURE 5:**
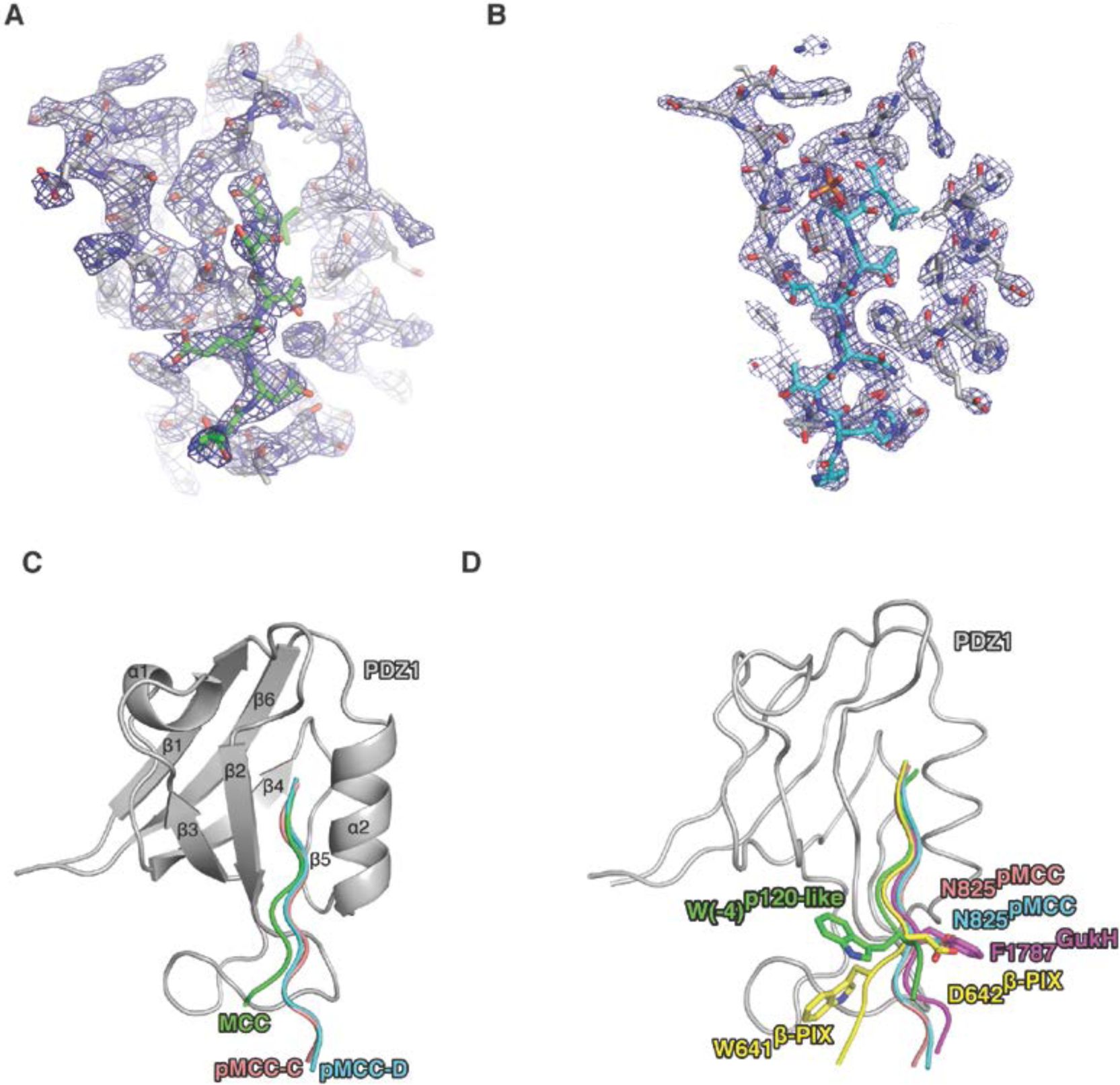
2Fo-Fc electron density maps of PDZ1 complexes with wild-type MCC and phosphorylated MCC peptide. (A) Electron density map encompassing the binding groove of Scribble PDZ1 in complex with MCC peptide. PDZ1 is shown as grey sticks whereas MCC peptide is shown as green sticks. The electron density map is shown as a blue mesh contoured at 1.5 σ. (B) Electron density map encompassing the binding groove of Scribble PDZ1 in complex with pMCC. PDZ1 is shown as grey sticks whereas pMCC is shown as cyan sticks. The electron density map is shown as a blue mesh contoured at 1.5 σ. (C) Overlay of ribbon traces of PDZ1 (grey) bound to MCC (green) or pMCC peptides (cyan and coral). (D) Superimposition of Scribble PDZ1 (grey) bound to pMCC (cyan and coral) complexes with β-PIX (yellow, PDB ID 5VWK), p120-like synthethic peptide (green, PDB ID 1N7T), and Gukh (magenta, PDB ID 5WOU) complexes.

In the PDZ1:MCC structure, the pocket accommodating L829^MCC^ is formed by PDZ1 L738, I740, I742, V797 and L800. In addition, PDZ1 forms several hydrogen bonds with MCC, including side chain mediated T827^MCC^:H793^PDZ1^ and N825^MCC^:S748^PDZ1^ (Figure 4,5) as well as the main chain contacts from L738^PDZ1^, G739^PDZ1^ and I740^PDZ1^ with the carboxyl group of L829^MCC^ and inter main chain contacts between T827^MCC^:I742^PDZ1^. The final hydrogen bond is formed by T824^MCC^:T749^PDZ1^. Examination of the complex of phosphorylated MCC (pMCC) with Scribble PDZ1 complex reveals two copies of PDZ1:pMCC in the asymmetric unit (denoted as PDZ1-A:pMCC-D and PDZ1-B:pMCC-C) which featured differences in the formation of hydrogen and ionic bonds. A comparison of the PDZ1:MCC complex with PDZ1-A:pMCC-D reveals only a single major difference in the formation of hydrogen or ionic bonds, where R762^PDZ1^ forms an ionic bond with the phosphate group on pS828^pMCC^ from a neighboring molecule. Furthermore the orientation of the N-terminus of pMCC-D is different, with two additional residue, H823 and P822 being resolved. A comparison of the PDZ1:MCC complex with PDZ1-B:pMCC-C reveals that R762^PDZ1^ also forms an ionic bond with the phosphate group on pS828^pMCC^, however in this instance the bond is formed with the actual pMCC chain bound in the PDZ1 domain binding groove. In addition, hydrogen bonds between S761^PDZ1^ and E826^pMCC^ as well as S748^PDZ1^ and N825^pMCC^ and G747^PDZ1^ and H823^pMCC^ are found. Interestingly, the T824^MCC^:T749^PDZ1^ hydrogen bond seen in the PDZ1:MCC complex is absent in the PDZ1:pMCC-C, and instead T749^PDZ1^ engages P822^pMCC-D^. Lastly, the C-terminal pMCC residue L829 only makes two contacts via its carboxyl group with PDZ1 via L738^PDZ1^ compared to the three contacts in PDZ1:MCC. The overall configuration of pMCC when bound to PDZ1 differs from wild-type MCC bound to PDZ1, with the C-terminus from both pMCC complexes revealing the position of H823 and P822 which are disordered in PDZ1:MCC (Figure 5 C).

Binding of a phosphorylated MCC peptide did not induce any significant overall structural changes to Scribble PDZ1 or the ligand binding groove compared to binding of unphosphorylated MCC, with the rmsd of a superimposition of the Scribble PDZ1:MCC complex with both pMCC complexes being 1.0 Å over 96 Cα atoms.

## DISCUSSION

Scribble is an important regulator of apico-basal cell polarity, and enables the integration of a diverse set of signalling pathways that all converge on Scribble via interactions using its four PDZ domains. An added layer of complexity to the interactions of Scribble with its interactors via PDZ domains is the possibility of post-translational modifications that alter the affinity with which Scribble binds to its partners. Indeed, several Scribble-ligand interactions have now been shown to be affected by post-translational modifications, suggesting that phosphorylation may be a major regulatory mechanism for Scribble-ligand interactions (14,21). Intriguingly, phosphorylation does not always impact Scribble PDZ domain binding. Phosphorylation of APC seems to have little effect on Scribble binding, whereas phosphorylation of RPS6KA2 leads to a 4-fold increase in affinity for Scribble PDZ1. In contrast, phosphorylation of MAPK12 completely abolishes Scribble PDZ1 interaction (14).

It has previously been shown that phosphorylation of MCC impacts binding to Scribble PDZ domains, with phospho-MCC showing a modest 2-fold increase in affinity for Scribble PDZ1 (14). Similarly, phospho-MCC has been predicted to show tighter binding to Scribble PDZ3 (21). However, NMR chemical shift analysis revealed no substantial differences for Scribble PDZ1 binding to phosphorylated and unphosphorylated MCC that could provide a structural rational for the reported increased affinity (23).

Our data suggests that whilst Scribble PDZ domains 1 and 3 are able to bind the PBM of MCC, the affinities were not significantly changed after phosphorylation of a Ser residue at the -1 position of the MCC PBM. These findings are compatible with the NMR chemical shift report but are unexpected, since previous reports indicated that phosphorylation of Ser at -1 of the MCC PBM is important for the interaction of MCC with Scribble and control of lamellipodia formation, and that loss of phosphorylation reduced the amount of MCC bound to Scribble in immunoprecipitation experiments. Our affinity measurements are broadly corroborated by crystal structures of Scribble PDZ1 domain bound to unphosphorylated and phosphorylated PBMs of MCC. Here, the phospho-Ser at position -1 in the pMCC PBM is involved in one major contact with R762 of Scribble PDZ1, as well as two additional hydrogen bond between S762^PDZ1^ and E826^pMCC^ as well as G747^PDZ1^ with H823^pMCC^. These interactions however are not uniform across both copies of the PDZ1:pMCC complexes present in the asymmetric unit, suggesting that the interplay between PDZ1 and pMCC and the required intermolecular interactions are flexible, thus providing a rationale for the modestly tighter binding of phospho-MCC to Scribble PDZ1 compared to unphosphorylated MCC. These findings are in agreement with findings from Sundell *et al.*, who based on chemical shift data speculated that phosphorylation at the -1 position of the MCC PBM may not be a major functional switch (23). Examination of the phosphorylated Ser at the -1 position of the MCC PBM bound to Scribble PDZ1 reveals that although pMCC is able to adopt an acidic clamp configuration with -1 (pS828) and -3 (E826), Scribble PDZ1 does not harbor a corresponding charged residue that could be targeted by the pMCC acidic clamp in a manner previously observed in the SNX27 PDZ in complex with 5-HT_4(a)_R or in complex with LRRC3B-pSer(-3), which features an acidic clamp formed by -3 and -5 PBM (15).

Although MCC is able to bind to both Scribble PDZ1 and 3 domains, the interaction with PDZ3 is substantially tighter. Interestingly, PDZ3 has been shown to be the primary binding site for MCC (16), with phosphorylation of S828 at the -1 position predicted to lead to tighter binding compared to unphosphorylated MCC (21) based on functional data where enzymatically dephosphorylated MCC showed reduced levels of immunoprecipitation with full length Scribble. Unexpectedly, our data suggest that phosphorylation of MCC -1 S828 does not significantly alter its affinity for Scribble PDZ3. Pangon *et al.* (21) previously observed that there is no significant difference in immunoprecipitation levels between unphosphorylated MCC and phosphorylated MCC with Scribble, whereas an MCC S828A mutant displayed reduced Scribble binding. These findings were interpreted as evidence that MCC is usually phosphorylated. Our data suggest that MCC may not necessarily be phosphorylated to account for the lack of significant differences in immuno-precipitation levels observed, and may be indicative of an effect that is not directly linked to Scribble binding. This is in contrast to MCC binding to Dlg, where phosphorylation of MCC leads to a decrease of affinity from 2.7 to 95 μM for Dlg PDZ2 (14).

Examination of the thermodynamics of MCC and pMCC binding to PDZ3 reveal that the tighter binding of pMCC to PDZ3 is due to more favorable-TΔS or entropy terms rather than enthalpy terms, which remained the same. We speculate that few or no major additional hydrogen bonds are formed with the phosphate group on Ser -1, however this would have to be confirmed by direct structure determination of a PDZ3:pMCC complex.

Compared to other previously established affinities of Scribble PDZ1 domain for interactors, MCC is not a particularly high affinity interactor. Gukh engages *Drosophila* Scribble PDZ1 with 660 nM affinity (26), and β-PIX binds human Scribble PDZ1 with 2 μM affinity (24,27). MCC only binds to Scribble PDZ1 with 7.7 μM affinity, and examination of the different molecular contacts between MCC and Scribble PDZ1 indicate that the lower affinity compared to β-PIX may be due to the lack of a hydrophobic residue engaging the PDZ1 β2-3 loop, as is achieved by β-PIX via β-PIX W641 (24) or Trp at position -4 in a high affinity nanomolar complex of a synthetic peptide with Erbin (28) (see Figure 5D). An alternative configuration that enables high affinity interactions is observed in the Scribble PDZ1:Gukh complex, where F1784 in Gukh contacts H796 on the opposite side of the ligand binding groove (26).

In summary, we show that Scribble PDZ3 domain binds MCC with a ∼2-fold higher affinity compared to Scribble PDZ1 domain. Furthermore, phosphorylation of MCC at the -1 position of the PBM only has a modest impact on binding to Scribble PDZ1 and PDZ3 domains, suggesting that the impact of phosphorylation of the MCC PBM at the -1 position extends beyond a simple change in binding affinity to Scribble PDZ domains and that could be dependent on a Scribble PDZ interdomain interaction to favour or block MCC binding. Another possibility is that -1 phosphorylation of the MCC PBM primarily affects an interactor that indirectly affects Scribble such as Dlg, which has been shown to display substantial modulation of affinity for MCC depending on the phosphorylation status of the MCC PBM.

In summary, our findings provide a clear structural basis for Scribble PDZ1:MCC interactions, and will enable more detailed structure-guided investigations to understand the precise effect of MCC phosphorylation on the control of cell polarity and directed migration.

## EXPERIMENTAL PROCEDURES

### Protein expression and purification

Protein expression constructs encoding the PDZ domains of human Scribble (Uniprot accession number: Q14160) PDZ1 (728–815); PDZ2 (833–965); PDZ3 (1005–1094); and PDZ4 (1099– 1203)) were obtained as synthetic cDNA codon optimized for *Escherichia coli* expression and cloned into the pGil-MBP (29) and pGex-6P3 (GE Healthcare). Protein overexpression was performed using *Escherichia coli* BL21 (DE3) pLysS cells (BIOLINE) in super broth supplemented with 200 µg/mL ampicillin (AMRESCO) using auto-induction media (10 mM Tris-Cl pH7.6, 100 mM NaCl, 1 mM MgSO4, 0.2 % (w/v) D-lactose, 0.05 % (w/v) glucose, 0.5 % (v/v) glycerol) (30) at 37°C until the optical density at 600 nm (OD600) reached 1.0 before transferring cultures to 20°C for 24 hours for protein expression. Bacterial cells were harvested by centrifugation and lysed in the presence of deoxyribonuclease I (Sigma-Aldrich) from bovine pancreas using TS Series 0.75 kw model cabinet (Constant Systems Ltd.) at 25 kPsi, Qsonica Q700 sonicator at amplitude 50 for 4 minutes process time on ice (5 seconds pulse-on time and 30 seconds pulse-off time). Lysates were clarified by centrifugation at 20,000 x g for 20 minutes using an Avanti® J-E (Beckman Coulter). The resulting supernatant was filtered using Millex-GP syringe filter unit 0.22 µM (Merck Millipore) prior to loading onto equilibrated columns for affinity purification.

Glutathione-S-transferase (GST) tagged recombinant PDZ1 protein was captured using glutathione sepharose 4B (GE Healthcare) in buffer A (50 mM Tris-Cl pH 8.0, 150 mM NaCl and 1 mM EDTA) and was cleaved on-column with HRV 3C protease to remove the GST tag. Cleaved protein was retrieved with buffer A and GST tagged protein was eluted with buffer A supplemented with 20 mM reduced L-glutathione.

Hexahistidine maltose binding protein (His-MBP) tagged recombinant proteins (PDZ2, PDZ3 and PDZ4) were purified using 5mL HisTrap HP columns (GE Healthcare) in buffer B (50 mM Tris-Cl pH 8, 300 mM NaCl) and washed with buffer B supplemented with 20 mM Imidazole before eluting in buffer B supplemented with 300 mM Imidazole. His-MBP tagged recombinant proteins were cleaved with TEV protease in buffer B supplemented with 0.5 mM EDTA and 1 mM DTT before being subjected to a second round of affinity chromatography to remove cleaved HisMBP tag and uncleaved fusion protein. All cleaved target proteins were subjected to size exclusion chromatography using the HiLoad 16/600 Superdex 75 (GE Healthcare) equilibrated in 25 mM Tris pH8.0, 150 mM NaCl, and eluted as a single peak.

### Isothermal titration calorimetry

Purified human Scribble PDZ domains were used in titration experiments against 8-mer peptides spanning the C-terminus of human MCC isoform-2 (Uniprot accession number: P23508; PHTNETSL), a modified MCC peptide (pMCC) bearing a phosphorylated Ser residue at position -1 (PHTNET(phoS)L), as well as MCC_SD (PHTNETDL) bearing Ser to Asp substitution on position -1 and MCC_SE (PHTNETEL) bearing a Ser to Glu substitution at position -1 to determine the effect of phosphorylation on MCC affinity for Scribble PDZ domains. PDZ1 concentration was quantitated at 280 nm absorbance (A280nm) using a NanoDrop 2000/2000c UV-Vis Spectrophotometer (Thermo Scientific). Since the concentration of PDZ2, PDZ3 and PDZ4 could not be determined by measurements of Abs280nm due to lack of useful aromatic amino acids, protein concentrations were calculated using the Scope method (31) by measuring absorbance at 205 and 280 nm using a NanoDrop 2000 spectrophotometer (Thermo Fisher Scientific).

Titrations were performed at 25°C with a stirring speed of 750 rpm using the MicroCalTM iTC200 System (GE Healthcare). A total of 20 injections with 2 µL each and a spacing of 180 seconds were titrated into the 200 µL protein sample (25 mM Tris pH 8.0, 150 mM NaCl), except for the first injection which was only 0.4 µL. Protein concentration of 75 µM against peptide concentration of 0.9 mM were used. Peptides were purchased from Mimotopes (Mulgrave, Australia). Raw thermograms were processed with MicroCal Origin® version 7.0 software (OriginLabTM Corporation) to obtain the binding parameters of each interaction. A synthetic pan-PDZ binding peptide referred to as superpeptide (RSWFETWV) was used as a positive control (26).

### Protein crystallisation, data collection and refinement

Complexes of PDZ1 with MCC and pMCC peptides were reconstituted by mixing protein and peptide at a 1:2 molar ratio. The dilute protein complexes were then concentrated to 15 and 23 mg/ml respectively, using a 3-kDa molecular mass cut-off centrifugal concentrator (Millipore), flash-cooled, and stored under liquid nitrogen. Crystallization trials were carried out using 96-well sitting-drop trays (Swissci) and vapor diffusion at 20 °C either in-house or at the CSIRO C3 Collaborative Crystallization Centre, Melbourne, Australia. 0.15-μl of protein-peptide complexes were mixed with 0.15 μl of various crystallization conditions using a Phoenix nanodispenser robot (Art Robbins). Commercially available screening kits (PACT Suite and JCSG-plus Screen) were used for the initial crystallization screening, with hit optimization performed using a 96-well plate at the CSIRO C3 Centre. Crystals of PDZ1 in complex with MCC peptide were obtained at 15 mg/ml in 25% (w/v) polyethylene glycol 1000, 0.15 M lithium sulfate monohydrate and 0.1 M phosphate/citrate pH 4.5. The PDZ1-MCC crystals were cryo-protected using 20% (w/v) glucose and flash-cooled at 100 K using liquid nitrogen. Plate shaped crystals were obtained belonging to space group I121. Crystals of PDZ1 in complex with pMCC peptide were obtained at 23 mg/ml in 0.1 M Phosphate/citrate 4.2 and 40 % (v/v) PEG 300. The PDZ1-pMCC crystals were cryo-protected using 30% (w/v) ethylene glycol and flash-cooled at 100 K using liquid nitrogen. Plate like crystals were obtained belonging to space group I4_1_ with twin law -h,k,-l (structure refinement was not possible at I4_1_22 as suggested by Pointless).

All diffraction data were collected on the MX2 beamline at the Australian Synchrotron using an Eiger 16M detector with an oscillation range of 0.1° per frame using a wavelength of 0.9537 Å. PDZ1:MCC diffraction data was integrated with Xia2 (32) and scaled using AIMLESS (33) while PDZ1-pMCC data was integrated with XDSme (34) to detector resolution. PDZ1-pMCC diffraction was anisotropic with resolution limits of 2.8 Å along the a* and b* orientation and 2.1 Å on the c* orientation. Therefore, the data was elliptically truncated and corrected using the Staraniso server (http://staraniso.globalphasing.org).

Both structures were solved by molecular replacement using Phaser (35) with the structure of human Scribble PDZ1 (PDB code 5VWC (24) or 2W4F) as a search model. The final TFZ and LLG values were 16.7 and 253 for PDZ1-MCC and 25.6 and 723 for PDZ1-pMCC, respectively. The solutions produced by Phaser was manually rebuilt over multiple cycles using Coot (36) and refined using PHENIX (37). Data collection and refinement statistics details are summarized in Table 2. MolProbity scores were obtained from the MolProbity web server (38). Coordinate files have been deposited in the Protein Data Bank under the accession codes 6MTU and 6MTV. All images were generated using the PyMOL Molecular Graphics System, Version 1.8 Schrödinger, LLC. All software was accessed using the SBGrid suite (39). All raw diffraction images were deposited on the SBGrid Data Bank (40) using their PDB accession code 6MTU and 6MTV.

## Acknowledgements

We thank staff at the MX beamlines, in particular Tom Caradoc-Davis, at the Australian Synchrotron for help with X-ray data collection, and the CSIRO C3 Collaborative Crystallization Centre for assistance with crystallization and the Comprehensive Proteomics Platform at La Trobe University for core instrument support. We thank the ACRF for their support of the Eiger MX detector at the Australian Synchrotron MX2 beamline.

## FOOTNOTES

This work was supported in whole or part by the National Health and Medical Research Council Australia (Project Grant APP1103871 to MK, POH; Senior Research Fellowship APP1079133 to POH), Australian Research Council (Fellowship FT130101349 to MK) and La Trobe University (Research focus area “Understanding Disease” project grant).

